# Autocatalytic chemical networks preceded proteins and RNA in evolution

**DOI:** 10.1101/693879

**Authors:** Joana C. Xavier, Wim Hordijk, Stuart Kauffman, Mike Steel, William F. Martin

## Abstract

Modern cells embody metabolic networks containing thousands of elements and form autocatalytic molecule sets that produce copies of themselves. How the first self-sustaining metabolic networks arose at life’ s origin is a major open question. Autocatalytic molecule sets smaller than metabolic networks were proposed as transitory intermediates at the origin of life, but evidence for their role in prebiotic evolution is lacking. Here we identify reflexively autocatalytic food-generated networks (RAFs)—self-sustaining networks that collectively catalyze all their reactions—embedded within microbial metabolism. RAFs in the metabolism of ancient anaerobic autotrophs that live from H_2_ and CO_2_ generate amino acids and bases, the monomeric components of protein and RNA, and acetyl-CoA, but amino acids and bases do not generate metabolic RAFs, indicating that small-molecule catalysis preceded polymers in biochemical evolution. RAFs uncover intermediate stages in the origin of metabolic networks, narrowing the gaps between early-Earth chemistry and life.

## Introduction

Cells are autocatalytic in that they require themselves for emergence. The origin of the first cells from the elements on the early Earth roughly 4 billion years ago (Baross, 2018; Betts et al., 2018; Varma et al., 2018; Tashiro et al., 2017) must have been stepwise. The nature of autocatalytic systems as intermediate states in that process is of interest. Autocatalytic molecule sets are simpler than cellular metabolism and produce copies of themselves if growth substrates for food and a source of chemical energy for thermodynamic thrust are provided (Fuchs, 2011; Goldford et al., 2017; Semenov et al., 2016). In theory, sets of organic molecules should be able to form autocatalytic systems (Dyson, 1982; Eigen and Schuster, 1977; Kauffman, 1971), which, if provided with a supply of starting’ food’ molecules, can emerge spontaneously and proliferate via constraints imposed by substrates, catalysts, or thermodynamics (Kauffman, 1986). Autocatalytic sets have attracted considerable interest as transitory intermediates between chemical systems and genetically encoded proteins at the origin of life (Hordijk et al., 2010; Kauffman, 1986; Smith and Morowitz, 2004; Sousa et al., 2015), but they have not been identified in non-enzymatic metabolic networks so far and evidence for their existence during prebiotic evolution is lacking.

Of special interest for metabolic evolution are a class of mathematical objects called Reflexively Autocatalytic Food-generated networks—RAFs—in which each reaction is catalyzed by a molecule from within the network and all molecules can be produced from a set of food molecules by the network itself (Hordijk and Steel, 2004). Small chemical systems resembling RAFs have been constructed in the laboratory (Ashkenasy et al., 2004; Semenov et al., 2016; Vaidya et al., 2012), although still far from the scale of cellular metabolism, which is composed of thousands of reactions. Modern cellular metabolism is enzyme–based, but >60% of enzyme mechanisms described to date involve one or more cofactors (Ribeiro et al., 2018) and 40% of all proteins crystallized have a bound metal relevant to their function (Guengerich, 2016). RAFs can thus be identified in modern metabolism (Sousa et al., 2015) by attributing the catalysis of enzymes to their metals and cofactors in prebiotic evolution (Argueta et al., 2015; Martin and Russel, 2007; Stockbridge et al., 2010; Varma et al., 2018; White, 1976; Zabinski and Toney, 2001). If autocatalytic chemical networks antedate genetically encoded proteins, cofactor-dependent RAFs might have been involved and, if so, should have left evidence for their existence in modern metabolic networks.

In search of RAFs, we investigated different levels of ancient metabolism preserved in modern cells. Starting with the biosphere level of the KEGG database, we first removed all eukaryote-specific reactions, and then peeled back one more layer of time by examining anaerobic metabolism. The detection of a large RAF in anaerobic prokaryotic metabolism prompted us to ask whether RAFs are also preserved in the metabolism of ancient anaerobic autotrophs that trace to the last universal common ancestor, LUCA (Weiss et al., 2016). As far back as we could look in metabolic evolution, RAFs were found. They were found in the metabolism of the acetogenic bacterium *Moorella thermoacetica* and the methanogenic archaeon *Methanococcus maripaludis*, primitive lineages that live on the simplest source of carbon and energy known, the H_2_–CO_2_ redox couple (Baross, 2018; Fuchs, 2011; Martin and Russel, 2007; McCollom and Seewald, 2007; Müller et al., 2018; Weiss et al., 2016). Their RAFs furthermore intersect in a primordial network that generates amino acids, nucleosides, and acetyl-CoA from a starting set of simple food molecules, shedding light on the nature of autocatalytic networks that existed before the first cells arose from the elements on the early Earth.

## Results

### Two-thirds of global prokaryotic metabolism can be annotated with small-molecule catalysis

In search of RAFs in 4-billion-year-old metabolism, we started from all 10,828 KEGG reactions and purged the set of non-primordial reactions in two pruning steps. First, we removed reactions assigned only to eukaryotes. Such reactions are not primordial because eukaryotes arose less than 2 billion years ago (Betts et al., 2018). Second, we excluded O_2_-dependent reactions, because O_2_ is a product of cyanobacterial photosynthesis, which arose about 2.4 billion years ago (Fischer et al 2016). These pruning steps left 5847 enzyme-associated reactions, 66% of which involve at least one cofactor. The addition of 147 spontaneous reactions generated a global network comprising 5994 reactions and 5723 metabolites (see Materials and Methods, **Figure S1 and Table S1A**). The cofactors involved in this ancient anaerobic network are distributed among the five different Enzyme Commission (E.C.) classes as shown in **Figure 1**. Metal catalysis is widespread across all classes of metabolism, and NADH dominates the oxido-reductase reactions. The network comprises 70% of the initial enzymatic reaction network before removal of O_2_-dependent and eukaryote-specific reactions, indicating that most metabolism was invented in the anaerobic world (Raymond and Segrè, 2006).

**Figure 1.**
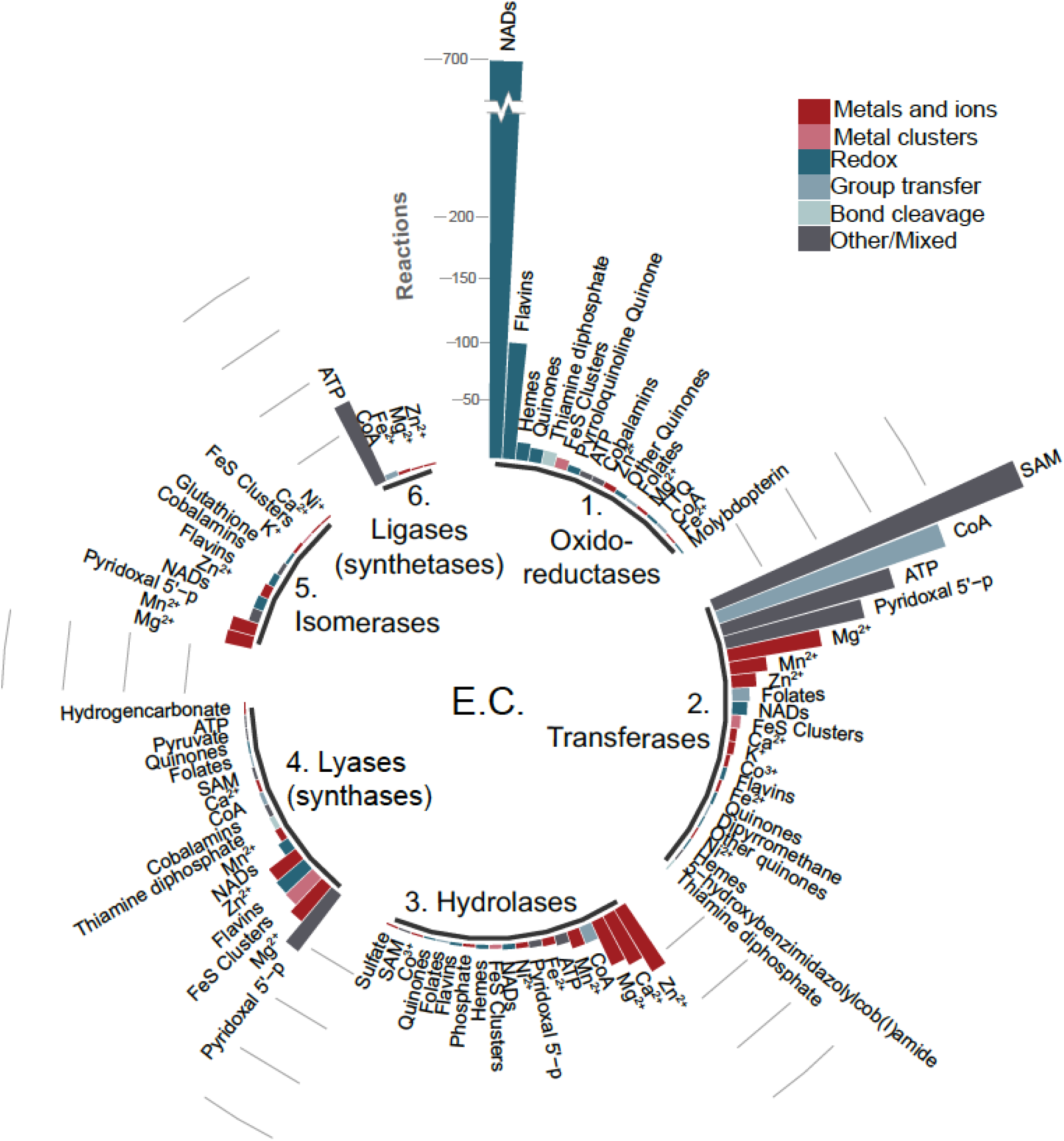
Catalysts in global oxygen-independent prokaryotic metabolism. The catalysis-annotated network separated by Enzyme Commission (EC) classes with the corresponding cofactors for each. Cofactors are grouped (legend, top–right) according to their function in catalysis.

### Autocatalysis in global metabolism expands with a small set of cofactors

The largest possible RAFs (maxRAFs) in a network are of interest because they represent its largest component of autocatalytic complexity. **Figure 2A** shows a schematic representation of a RAF within a metabolic network. The maxRAFs in the global prokaryotic O_2_-independent network were identified for different food sets, that is, molecules provided by the 3-environment (**Table S2**). An inorganic food set containing H_2_O, H_2_, H^+^, CO_2_, CO, PO_4_^3−^, SO_4_^2−^, HCO_3_^−^, P_2_O_7_^4−^, S, H_2_S, NH_3_, N_2_, all metals, FeS clusters and other metal clusters, a generalist acceptor, donor, and metal produced a minute maxRAF with eight reactions linking ammonia, carbon, and sulfide transformations. The addition of formate, methanol, acetate, and pyruvate, which are central metabolites with experimental evidence for synthesis from CO_2_ and metals (Varma et al., 2018), doubles the maxRAF size to 16 reactions. In principle, the addition of organic cofactors (**Table S2**) to the food set should generate larger maxRAFs. Sequential addition of the eight most frequent cofactors identified in the last universal common ancestor’s (LUCA’ s) proteins (Weiss et al., 2016) to the metal–CO_2_ food set expanded the maxRAF from 16 to 914 reactions (**Figure 2B**). Addition of all cofactors germane to the anaerobic network generates a maxRAF with 1335 reactions spanning 25% of the starting anaerobic network. Sequential addition of the eight compounds that were most frequent in that maxRAF, to the metal–CO_2_ food set, expands the maxRAF from 16 to 1066 reactions, whereas sequential addition of the five compounds with the greatest impact (upon removal from the food set) on anaerobic maxRAF size followed by the three most frequent in the largest maxRAF yields a final maxRAF of 1248 reactions (**Figure 2B**). These results indicate that RAFs can grow in size through sequential incorporation of organic cofactors (**Figure 2B**). RAFs can thus provide structure, contingency, increasing complexity, and direction to interactions among molecule food sets, given a sustained geochemical source of carbon, energy, and electrons.

**Figure 2.**
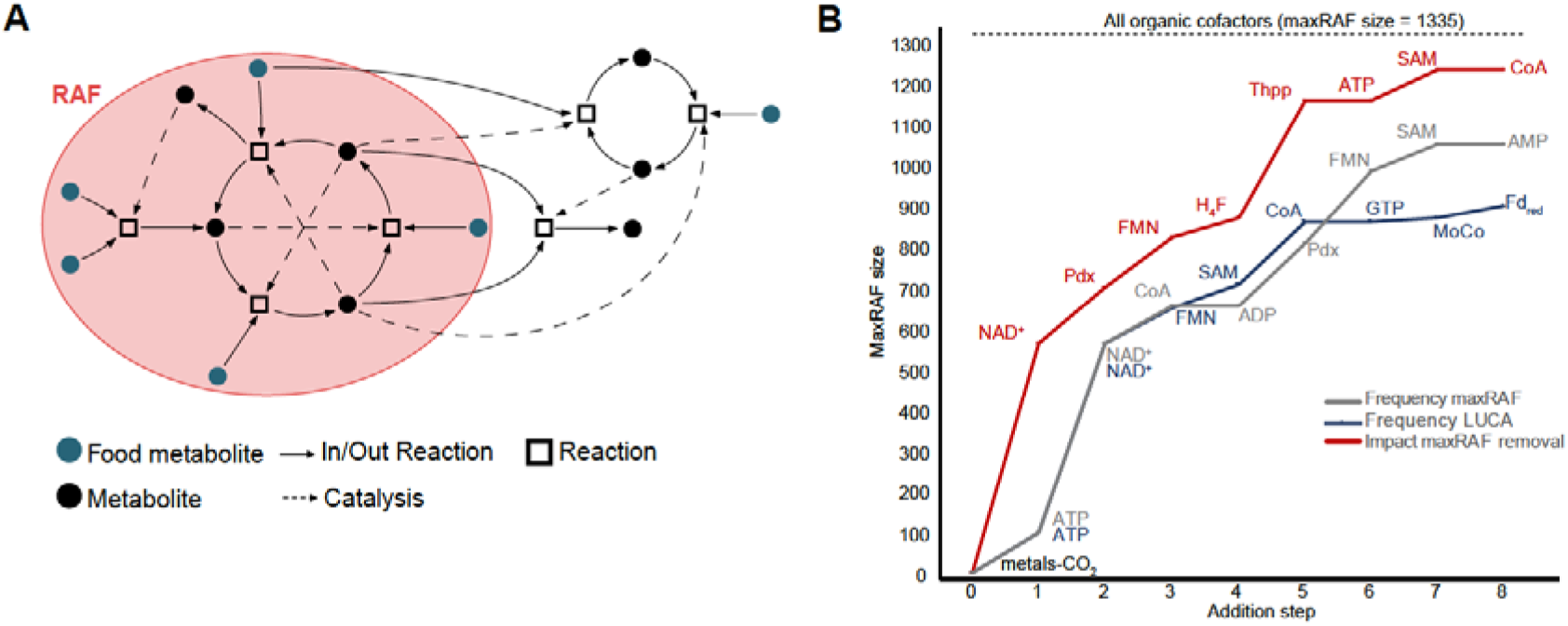
Autocatalysis in global metabolism expands with a small set of cofactors. **(A)** Schematic depiction of a reflexively autocatalytic food-generated network (RAF) highlighted (red ellipse) in a metabolic network. Food metabolites (green circles) may enter the RAF allowing subsequent reactions (squares) to occur and other metabolites (black circles) to be produced. Each reaction is catalyzed by a metabolite in the network (catalysis shown in dashed arrows). **(B)** Increasing maxRAF sizes with the sequential addition to the food set of the organic cofactors (i) with the highest impact on maxRAF size upon removal (red) (ii) most frequent in the maxRAF with all organic cofactors added (grey), and (iii) most frequent in enzymes predicted to be in LUCA (Weiss et al., 2016) (blue) (Pdx – pyridoxal 5-phosphate; H_4_F – tetrahydrofolate, Thi – thiamine diphosphate, MoCo – molybdopterin; Fd_red_– reduced ferredoxin). Top dashed line shows the maxRAF size obtained when all organic cofactors are added to the food set.

### Autocatalytic networks point to an early autotrophic metabolism

If autocatalytic sets were instrumental at the origin of metabolism (Kauffman, 1986), lineages with a physiology very similar to that of the first cells should harbor the most ancient RAFs. Several lines of evidence indicate that methanogens and acetogens reflect the ancestral state of microbial physiology: they live on the simplest source of carbon and energy known, the H_2_–CO_2_ redox couple (Baross, 2018; Fuchs, 2011; Martin and Russell, 2007; McCollom and Seewald, 2007; Müller et al., 2018; Weiss et al., 2016), they assimilate geochemically-generated carbon species (Stupperich and Fuchs 1984; Lang et al. 2010), they generate ATP from CO_2_ fixation (Fuchs, 2011), their core bioenergetic reactions occur abiotically in hydrothermal vents (McCollom and Seewald, 2007; McDermott et al., 2015) and under laboratory conditions (Varma et al., 2018), their ecology and gene trees link them to LUCA (Weiss et al., 2016), and they still inhabit primordial habitats within the crust today (Ijiri et al. 2018). Subsets of the global prokaryotic O_2_-independent network were obtained by parsing the genomes of the acetogen *Moorella thermoacetica* (Ace) and the methanogen *Methanococcus maripaludis* (Met). These were completed with reactions from corresponding manually curated genome-scale metabolic models (Islam et al., 2015; Richards et al., 2016), resulting in 1193 reactions for Ace and 920 for Met (**Tables S1B and S1C**). Both the acetogen and the methanogen metabolic networks contain RAFs. When all organic cofactors are added to the food set, the maxRAFs contain 394 and 209 reactions for Ace and Met, respectively, spanning major KEGG functional categories (**Figure 3; Tables S2 and S3, Figures S2 and S3**).

**Figure 3.**
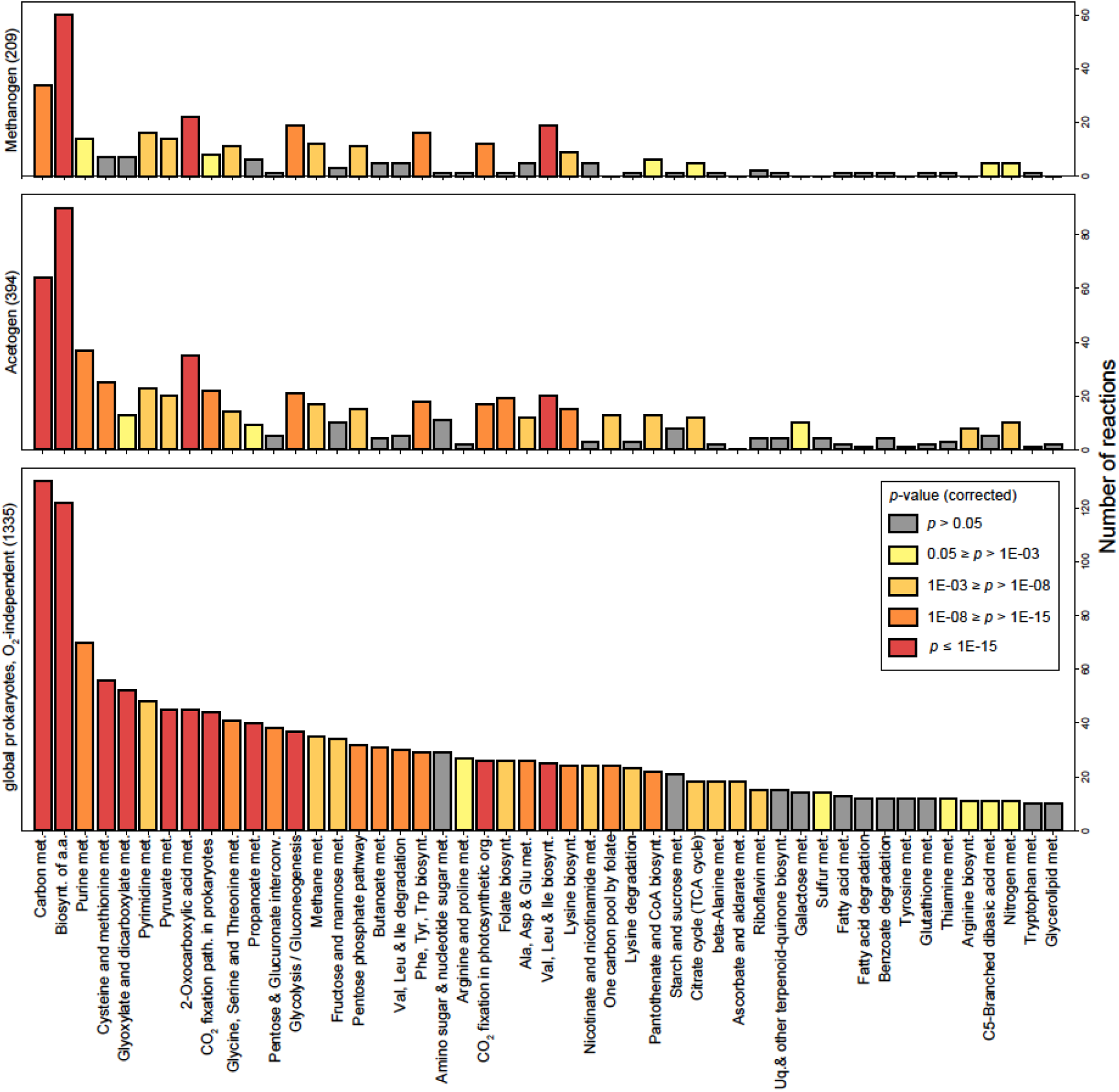
Autocatalytic networks point to an early autotrophic metabolism. Number of reactions in each functional category for three maxRAFs and functional enrichment compared with the global O_2_-independent prokaryotic network. Colors represents bins of corrected *p*-values (Fisher’s exact test with Benjamini–Hochberg FDR multiple-testing correction). From bottom to top, maxRAF obtained for (sizes in brackets): global O_2_-independent prokaryotic network, acetogen (Ace) and methanogen (Met). Categories are sorted according to the number of reactions in the first maxRAF, from smallest to largest; only categories where this maxRAF had more than 10 reactions are shown.

Carbon fixation and biosynthetic pathways are represented and amino acids biosynthesis is highly enriched in all maxRAFs, recovering autotrophic components of early autocatalytic metabolism. The addition of peptide catalysis increases the maxRAF sizes obtained with the global anaerobic network, Met, and Ace by 93%, 47%, and 25% respectively (**Table S2**). This indicates that adding protein catalysis expands cofactor-supported autocatalytic sets, but does so to a much lesser degree in the metabolism of Met and Ace than it does in the global O_2_-independent prokaryotic network.

### LUCA’s metabolism was autocatalytic and autotrophic

The intersection of the Ace and Met maxRAFs should be more ancient than either. Three-quarters of the (smaller) Met maxRAF overlap with the (larger) Ace maxRAF in a connected network harboring 172 reactions and 175 metabolites (**Figures 4 and 5**; individual maxRAFs from Ace and Met in **Figures S2 and S3**). Six metabolites are disconnected, meaning the species interconvert them using different pathways; one example is that of glucose, catabolism of which arose after LUCA (Schönheit et al., 2016). Highly connected food metabolites in the primordial network (more than 13 edges) include H_2_O, ATP, protons, phosphate, CO_2_, NAD^+^, pyruvate, ammonia, diphosphate, coenzyme A and AMP; highly connected non-food metabolites (more than eight edges) include ADP, NADH, and other pyridine dinucleotides, glyceraldehyde-3-phosphate, and acetyl-CoA (**Table S4**). The network is able to produce six amino acids—asparagine, aspartate, alanine, glycine, cysteine, and threonine—plus the two nucleosides UTP and CTP. Cytochromes and quinones do not figure into the network.

**Figure 4.**
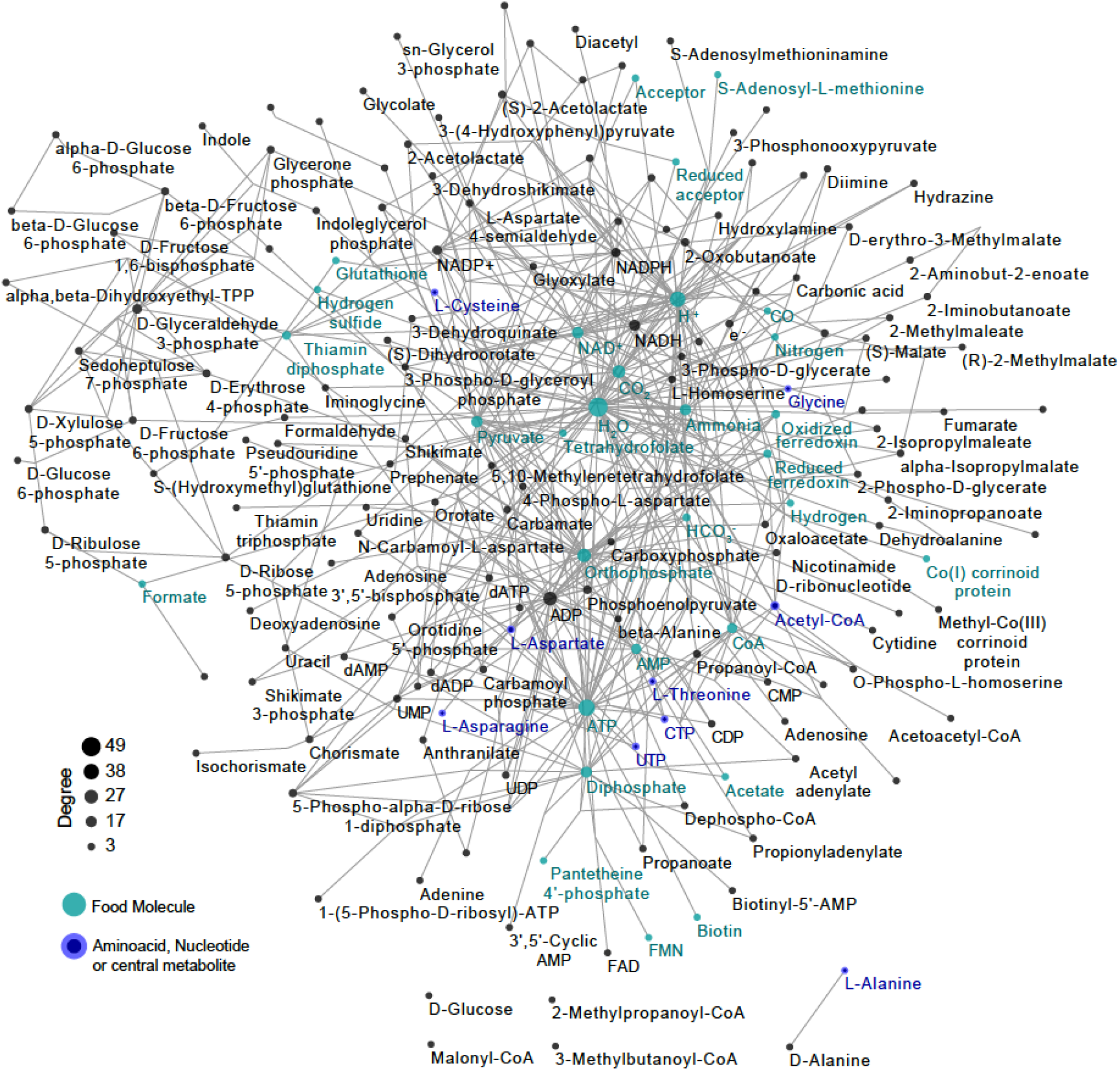
Core autocatalytic metabolism of the last universal common ancestor (LUCA). Intersection of the maxRAFs obtained with the networks of *Moorella thermoacetica* and *Methanococcus maripaludis* with a food set with organic cofactors (only metabolic interconversions are depicted; catalysis arcs are omitted for clarity). Six metabolites, including D-glucose and L-alanine (bottom) are in the intersection but disconnected from the remaining network. The node size is scaled according to the degree, with food molecules highlighted in green and relevant products in dark blue. ‘Acceptor’ and ‘Reduced Acceptor’ are abstract redox molecules as represented in KEGG metabolism.

**Figure 5.**
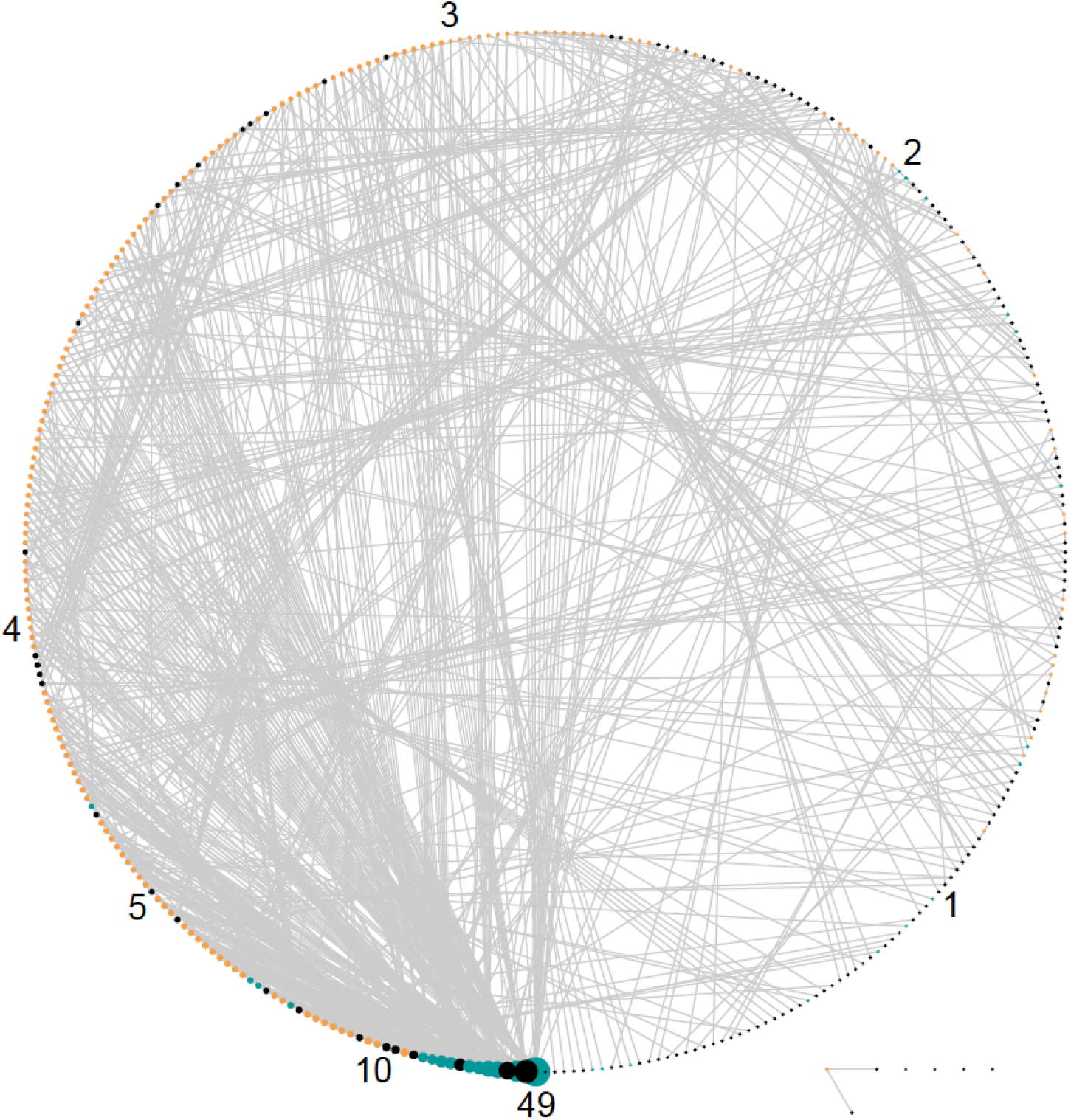
Circular bipartite representation of the autocatalytic metabolism in LUCA. Reactions (in orange) and metabolites (in green if food, grey for the rest) are represented as nodes. Nodes are sorted according to degree clockwise starting from the bottom; numbers show the degree at the respective position. ATP, the second most connected metabolite, can be removed from the food set without impact, therefore here is represented in black.

A different look at the primordial network reveals a hierarchical and highly-connected organization (**Figure 5**). The network is structured with a core half-moon, where the degree varies from 49 to 4 (**Table S4**). Food molecules cluster in the most connected area, showing the spark of autocatalytic metabolism by a handful of substrate molecules with degree higher than 10.

### LUCA’s metabolism is enriched in metal catalysis, ancient genes and autotrophic functions

In search of the distinct contributions for autocatalysis, we tested for enrichment in individual catalysts, functions and ancient genes encoding for reactions in the primordial network. There is a significant enrichment for metal and metal–sulfur cluster catalysis (**Figure 6A**), whereas thiamine diphosphate (a carrier of C2 units in metabolism) is the only organic cofactor that is significantly enriched in catalyzing the primordial network when compared with the global network, even though several others are present and essential for the network to grow (**Figure 6B**). The primordial network is also enriched in reactions for amino acid biosynthesis, carbon metabolism, and 2-oxocarboxylic acid metabolism when compared with the global network (**Figure 6B**). Comparing reactions in the primordial network to those catalyzed by genes that can be traced to LUCA by independent phylogenetic criteria (Weiss et al., 2016) uncovers highly significant enrichment relative to both the global network and its maxRAF (**Figure 6C**). The maxRAF obtained within the primordial network contains 120 reactions and is enriched in amino acid and carbon metabolism but produces cysteine as the sole amino acid, which is noteworthy because cysteine is the hub of sulfur metabolism and also is the sole ligand for incorporating Fe–S and Fe–Ni–S clusters in proteins (**Figure S4**).

**Figure 6.**
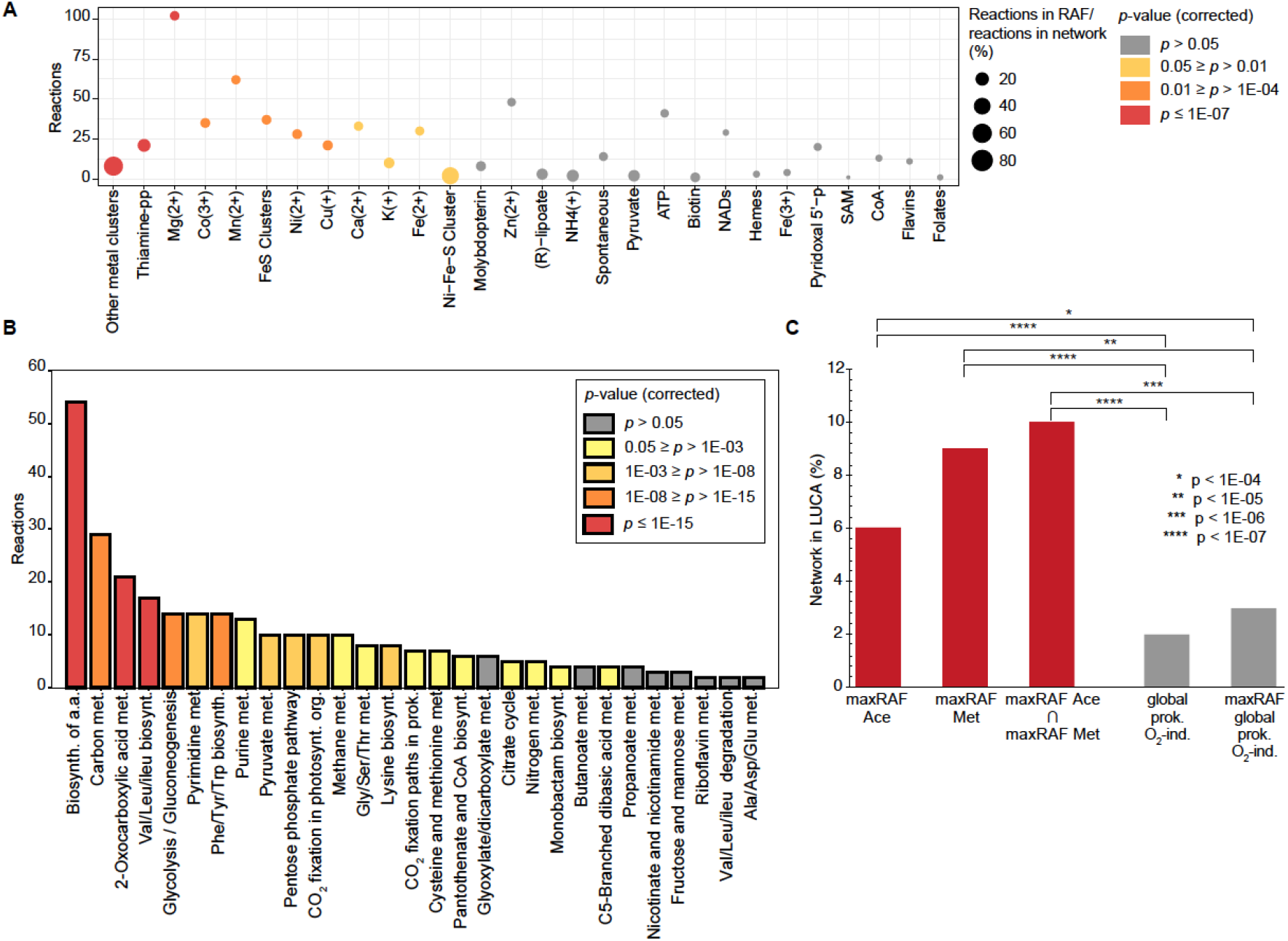
Properties of the core autocatalytic metabolism of the last universal common ancestor (LUCA). **(A)** Enrichment of cofactors catalyzing the reactions in the overlapping network between the acetogen and methanogen maxRAFs compared with the global O_2_-independent prokaryotic network. Circle size indicates the ratio between reactions in the intersection network and reactions in the global network; color indicates the corrected *p*-value (Fisher’s exact test with Benjamini–Hochberg FDR correction). **(B)** Enrichment of KEGG pathways in the overlapping network compared with the global O_2_-independent prokaryotic network. Color indicates bins of corrected *p*-values (Fisher’s exact test with Benjamini–Hochberg FDR correction). **c.** Proportion of metabolic networks predicted to be in LUCA (Weiss et al., 2016) and enrichment of the maxRAFs (red) compared with the global network and the maxRAF obtained with it (grey) (Fisher’s exact test).

### Autocatalysis before ATP and polymers

Crucial catalysts can be identified by removing them from the food set. NAD^+^ is strongly embedded in the RAF and its removal reduces the size of the maxRAF by ~50% (**Figure 7**). Other compounds heavily impacting maxRAF size are Fe–S clusters, pyridoxal-5-phosphate and divalent metals. Surprisingly, when we remove ATP from the food set of organic cofactors, this has no impact on the size of the maxRAF, both for the individual networks (**Figure 7**) and LUCA’s network (**Figure 5**). Why does ATP removal have such a small effect on RAFs? The simplest explanation is that ATP was not the primordial energetic currency (Goldford et al., 2017). This points to the increasingly evident role of energy currencies other than ATP in primordial metabolism, such as acyl phosphates (Martin and Russell, 2007), thioesters (Semenov et al., 2016), and reduced ferredoxin (Herrmann et al., 2008, Müller et al., 2018). Alternative energy currencies are particularly common in anaerobes (Müller et al., 2018).

**Figure 7.**
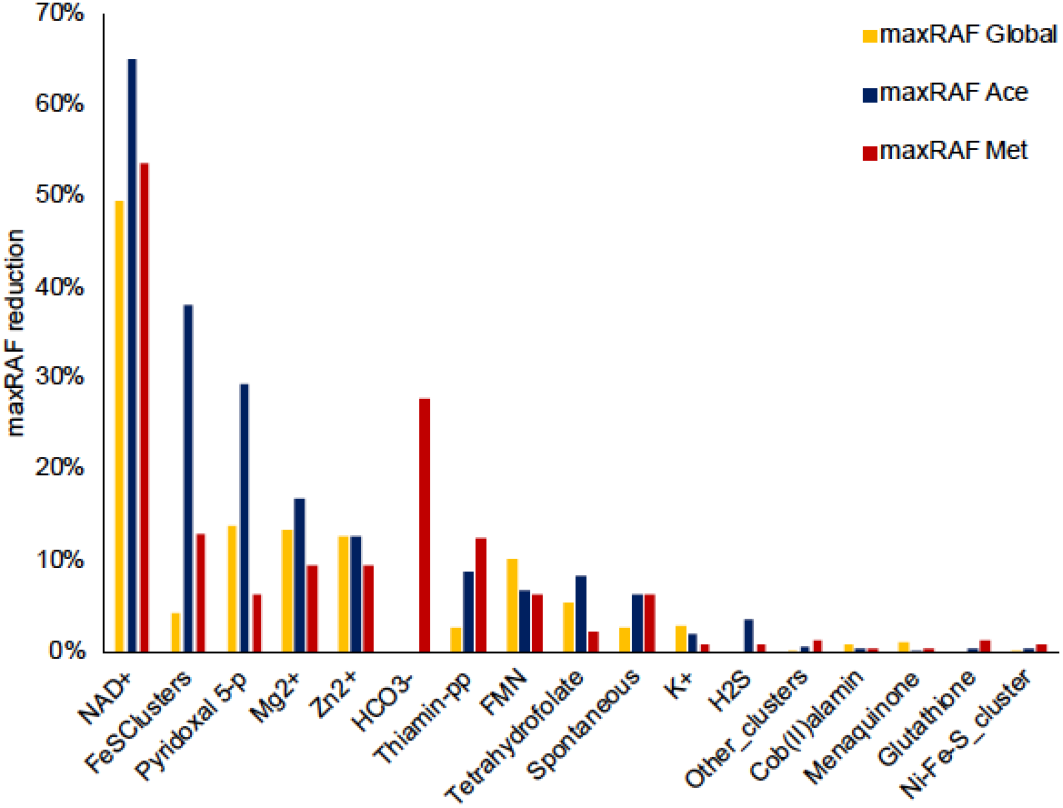
Impact of removing single molecules from the food set with organic cofactors on the size of maxRAFs. The impact is shown as the reduction in size of the maxRAF (percentage of the initial network lost) when each molecule is removed from the food set with all organic cofactors, for the global prokaryotic O_2_-independent network (yellow), *Moorella thermoacetica* (dark blue) and, *Methanococcus maripaludis* (red).

RAFs provided with a food set containing catalysts can generate amino acids and bases (**Figure 4**), but the converse is not true: adding amino acids and bases to the simplest food set, which includes inorganic catalysts and CO_2_ (**Table S2**), produces a miniscule 33-reaction maxRAF (**Figure S5**). The maxRAF contains 47 metabolites, 27 of which are food molecules. This indicates that autocatalytic networks embedded in microbial metabolism generated amino acids and bases using small-molecule catalysis prior to the advent of nucleic acids or peptide polymers.

## Discussion

Autocatalytic networks are objects of molecular self-organization (Dyson, 1982; Eigen and Schuster, 1977; Kauffman, 1986). Their salient property in the study of early biochemical evolution is the capacity to grow in size and complexity. Compounds generated from the food set become part of the network, hence autocatalytic networks can start small and grow, in principle to a size approaching the complexity of metabolic networks of modern cells (Hordijk et al., 2010), and very little catalysis by individual elements is required for autocatalytic networks to emerge (Hordijk and Steel, 2004; Mossel and Steel, 2005). Reflexively autocatalytic and food-generated networks—RAFs—are a particularly interesting formalization of collectively autocatalytic sets, as they capture a property germane to life: they require a constant supply of an environmentally provided food source in order to grow (Hordijk and Steel, 2004). In that sense, RAFs reflect metabolic networks in real cells, in that growth substrates are converted to end products, a proportion of which comprises the substance of cells. But RAFs are far simpler than metabolism because they can start very small.

RAFs have not been applied in the study of the evolution of chemical networks that led to the metabolism of modern cells, themselves large natural autocatalytic networks. By embracing the simple and robust premise that reactions catalyzed by simple molecules and inorganic compounds preceded metabolic reactions catalyzed by enzymes (Sousa et al., 2015; Stockbridge et al., 2010; White, 1976), we have retooled RAFs into an analytical instrument to investigate the nature of metabolic evolution.

Our analyses start with the enzymatic and spontaneous reactions charted in modern metabolism and use RAFs as a filter to uncover elements with self-organizational properties, to address the nature of processes in the earliest phases of evolution, before the origin of eukaryotes and before the appearance of oxygen. We find evidence for a role of autocatalytic networks in the early evolution of metabolism. The largest RAF that we identified in the whole prokaryotic anaerobic biochemical space has 1335 reactions and points to early autotrophy. This RAF is larger than the genome size of the smallest free-living archaeon, *Methanothermus fervidus* (Martínez-Cano et al., 2015). With a genome coding for 1,311 proteins and 50 RNA genes, *M. fervidus* lives from H_2_ and CO_2_ as carbon and energy sources (the food set) and requires only inorganic, geochemical nutrients, no other cells for survival (Anderson et al., 2010). H_2_ and CO_2_ were present in abundance on the early Earth and may have given rise to the first metabolic pathways that brought forth the first archaeal and bacteria cells (Goldford et al., 2017, Varma et al, 2018; Weiss et al., 2006). Our anaerobic RAF is however smaller than the reaction network in the smallest genome of bacteria that live from H_2_ and CO_2_, which is found in the acetogen *Thermoanaerobacter kivui*, encoding 2,378 proteins (Hess et al., 2014).

*M. fervidus* and *T. kivui* harbor primitive forms of methanogenesis and acetogenesis in that they both lack cytochromes and quinones, suggesting that they represent energy metabolic relics from the earliest phases of biochemical evolution on the primordial earth, before anaerobic respiratory chains had evolved (Martin and Russell, 2007). To investigate this aspect further, we examined the best annotated metabolic networks existing for H_2_-CO_2_ dependent archaea and bacteria, the methanogen *Methanococcus maripaludis* and the acetogen *Moorella thermoacetica*. Remarkably, a food set containing only small abiogenic molecules and a handful of organic cofactors generates sizeable RAFs in each of the networks, with 209 and 394 reactions respectively. The inclusion of organic molecules as catalysts in our food set is in line with a premise common to all scientific theories for the origin of life, namely that the environment provided starting material from which metabolism and life evolved.

RAFs uncover elements of metabolic evolution that go even further back in time before the divergence of archaea and bacteria from the last universal common ancestor, LUCA. The intersection of the RAFs of *M. maripaludis* and *M. thermoacetica* uncovers a fair amount in common—a core, conserved autocatalytic network with 172 reactions that is enriched in metal catalysis and carbon metal bonds (Martin, 2019) and also points both to early autotrophy and to the genes of LUCA (Weiss et al., 2016). Our results also show that the kickstart of autocatalysis in anaerobic metabolism does not require ATP. This is in accordance with the use of alternative energetic currencies in anaerobic prokaryotes (Müller et al., 2018) and recent findings that suggest that complexity in early metabolic reaction systems could have emerged without phosphate (Goldford et al., 2017).

An important insight uncovered by RAFs is the observation that although a food set with organic cofactors sparks a large autocatalytic network that generates amino acids and bases, the opposite does not occur: adding amino acids and bases to the simplest food set (which includes inorganic catalysts and CO_2_) only produces a minute RAF with 33 reactions. This result indicates that autocatalytic networks could generate amino acids and bases using catalysts prior to the advent of complex nucleic acids or peptide polymers. This stands in accordance with recent reports of amino acid synthesis catalyzed by native metals (Muchowska et al., 2019), and also with the physiology of extant anaerobic autotrophs: amino acids and bases are sequestered end-products of H_2_ and CO_2_ dependent metabolism, they are polymerized to make the substance of cells.

RAFs as a tool to study metabolic evolution can serve as a guide for the identification and construction of larger, biologically relevant autocatalytic reaction networks. The synthesis of compounds characteristic of the metabolism of acetogens and methanogens, intermediates and end products of the acetyl-CoA pathway and of the incomplete citric acid cycle from CO_2_ using only the catalysis of native metals (Muchowska et al., 2019), as well as the demonstrated catalytic power of organic cofactors without their enzymes including flavins (Argueta et al., 2015), pyridoxal 5’-phoshpate (Zabinski and Toney, 2001), SAM (Barrows and Magee, 1982) and NAD (Betanzos-Lara et al., 2012) encourages the investigation of more complex autocatalytic networks in laboratory reactors.

Our results are directly relevant to two deeply divided schools of thought concerning the nature of chemical reactions at the origin of life: genetics first and metabolism first. The genetics first school, or RNA world, holds that the origin of RNA molecules marked the origin of life-like processes, and that RNA both self-replicated and possessed catalytic abilities that led to the emergence of biochemical reactions (Orgel, 2008; Patel et al., 2015). In that view, the origin of the bases that drove that process forward is decoupled from biochemical processes that are germane to modern cellular metabolism. The metabolism-first school holds that spontaneous (exergonic) chemical reactions preceded reactions catalyzed by genetic material, and that those exergonic reactions continuously gave rise to substrate-product relationships (Martin and Russell, 2007; Morowitz et al., 2000). From such reactions, more complex interaction networks with autocatalytic properties arose (Kauffman, 2011; Smith and Morowitz, 2004), in which elements of the set intervened in reactions of the set, providing structure and direction to product accumulation. Our results show that in RAFs of anaerobic metabolism, nucleoside-related cofactors play a central role, albeit these have functional moieties that do not occur in RNA as it catalyzes protein synthesis. More importantly, our findings indicate that RNA could arise from metabolism, and the nature of the products accumulated in RAFs will include nucleic acids. In other words, RAFs applied to ancient autotrophic metabolism reveal a vector of autopoietic genesis that detects RNA emerging from metabolism rather than vice versa.

## Materials and Methods

### Catalysis-annotated metabolic networks

All the reactions and the EC numbers they are linked to were retrieved from KEGG (Kanehisa et al., 2017), along with their corresponding taxonomic annotations using the KEGG REST API (https://www.kegg.jp/kegg/rest/keggapi.html, accessed February 2018). The EC–reaction pairs were filtered by excluding reactions annotated only in eukaryotes. The corresponding chemical equations were then parsed to discard reactions involving molecular oxygen. Spontaneous reactions were parsed out of KEGG and added to the network with a fictional catalyst named “Spontaneous”. Reactions catalyzed by enzymes that are not spontaneous and the enzymes of which do not use any cofactors were assigned the catalyst “Peptide”.

Reactions that equate synonymous cofactors were added with the generic catalyst “Pooling”. Extensive curation was performed regarding catalysis rules, reaction reversibility, and amino acid production. The reversibility of reactions was parsed out of KGML files for KEGG pathways and manually–curated. The resulting set of reactions was then integrated with cofactor information from Uniprot (The Uniprot Consortium, 2018) through the corresponding EC numbers. Eighty-one unique cofactors were retrieved from Uniprot, which translated to 48 KEGG compounds or pools of catalytically equivalent cofactors linked to KEGG reactions through the EC numbers. Cofactors directly participating in reactions (NADs, ATP, SAM, CoA, Cobalamins, Folates, Flavins and Quinones) were extracted from the reaction stoichiometry if not assigned as cofactors in Uniprot. Of all EC numbers searched in Uniprot, 34% had at least one associated cofactor, 579 of which were EC numbers that involved more than one cofactor when parsed in a Boolean manner. All rules were added to the network as additional parameters. The subsets for Met and Ace were obtained by crossing the genomic annotation of *Moorella thermoacetica* and *Methanococcus maripaludis* in KEGG with the previously built network, and with the addition of missing reactions that were present in corresponding manually–curated models (Islam et al., 2015, Richards et al., 2016). The pipeline for the full procedure is shown in **Figure S1**.

### Detection of maxRAFs

All networks described above were tested for whether they contained maxRAFs with different food sets, which are described in the main text and available in **Table S2**. The fictional catalysts “Spontaneous” and “Pooling” were added in all tests, allowing for spontaneous reactions to always occur and synonymous cofactors to be equated. Pooling reactions that were part of the maxRAF were not accounted for in maxRAF sizes.

### RAF sets

We define a *chemical reaction system* (CRS) as a tuple *Q* = {*X, R, C, F*}, where:

- *X* = {*x*_1_, *x*_2_, …, *x*_*n*_} is a set of molecule types.
- *R* = {*r*_1_, *r*_2_, …, *r*_*m*_} is a set of reactions. A reaction *r* is an ordered pair *r* = (*A, B*) where *A, B* ⊂ *X*. The (multi)set *A* = {*a*_1_, …, *a*_*s*_} indicates the reactants and the (multi)set *B* = {*b*_1_, …, *b*_*t*_} indicates the products.
- *C* ⊆ *X* × *R* is a set of catalysis assignments. A catalysis assignment is a pair (*x*, *r*) with *x* ∈ *X* and *r* ∈ *R*, denoting that molecule type *x* can catalyse reaction *r*.
- *F* ⊂ *X* is a food set (i.e., molecule types that can be assumed to be available from the environment).

Given a CRS *Q*, a subset *R*′ of *R*, and a subset *X*′ of *X*, we define the *closure* of *X*′ relative to *R*′, denoted *cl*_*R*′_ (*X*′), to be the (unique) minimal subset *W* of *X* that contains *X*′ and that satisfies the condition that, for each reaction *r* = (*A, B*) in *R*′,

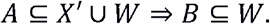

Informally, *cl*_*R*′_ (*X*′) is *X*′ together with all molecules that can be constructed from *X*′ by the repeated application of reactions from *R*′.

Given a CRS *Q* = {*X, R, C, F*} and a subset *R*′ of *R*, *R*′ is a *RAF set* if for each *r* = (*A, B*) ∈ *R*′

1. (Reflexive Autocatalysis): ∃*x* ∈ *cl*_*R*′_ (*F*):(*x, r*) ∈ *C*, and
2. (Food-generated): *A* ⊆ *cl*_*R*′_ (*F*).

In other words, a subset of reactions *R*′ is a RAF set if, for each of its reactions, at least one catalyst and all the reactants are in the closure of the food set relative to *R*′ (Hordijk and Steel, 2004).

### RAF algorithms

Given a CRS *Q* = {*X, R, C, F*}, an efficient (polynomial-time) algorithm exists for deciding whether *Q* contains a RAF set or not. It is presented formally in Algorithm 1.

#### Algorithm 1 RAF (*X, R, C, F*)

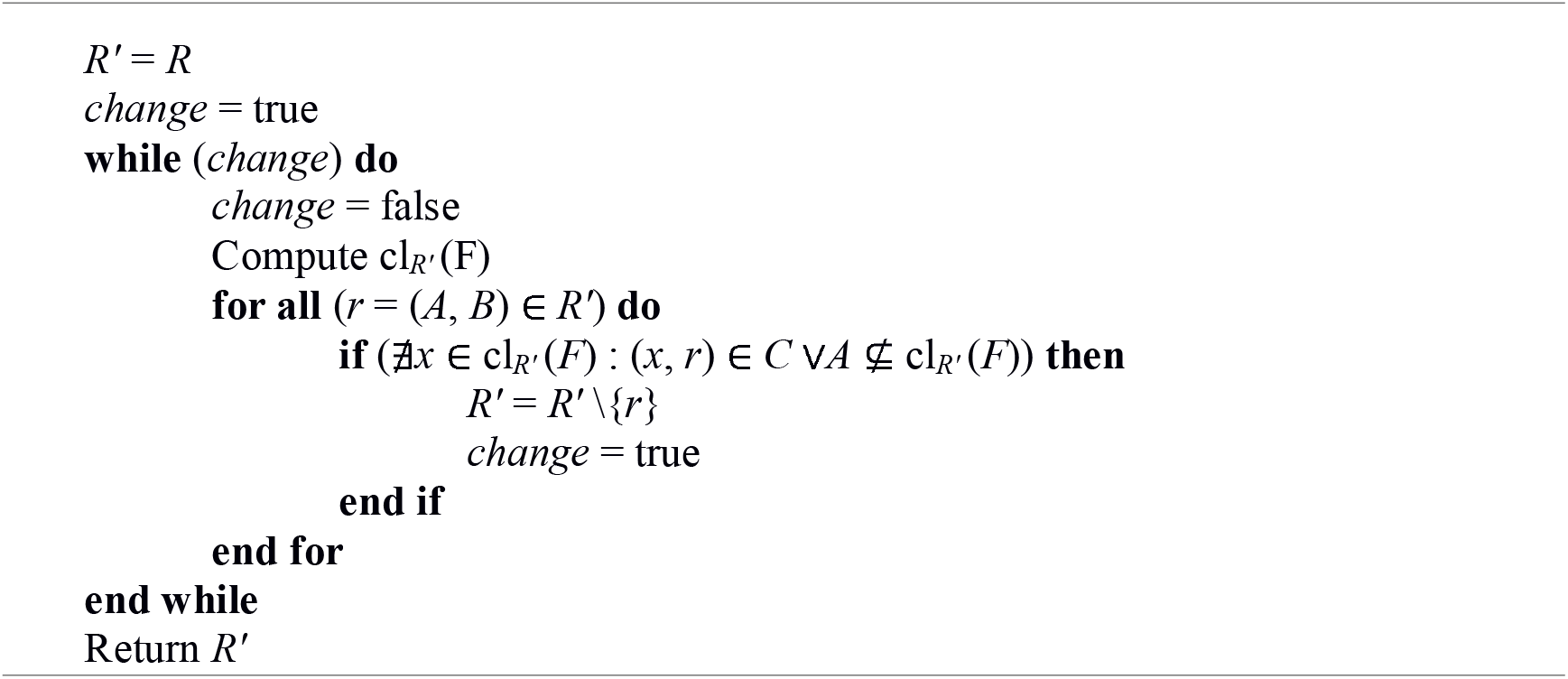

In plain words, starting with the full set of reactions *R*, the algorithm repeatedly calculates the closure of the food set relative to the current reaction set *R*′ and then removes all reactions from *R*′ that have none of their catalysts or not all of their reactants in this closure. This is repeated until no more reactions can be removed. If, upon termination of the algorithm, *R*′ is non-empty, then *R*′ is the unique *maximal* RAF set (maxRAF) contained in *Q* (i.e., a RAF that contains every other RAF in *Q* as a subset) (Hordijk and Steel, 2004). If *R*′ is empty, then *Q* does not contain a RAF set.

Computing the closure of the food set relative to the current reaction set *R*′ is computationally the most expensive step in the RAF algorithm. It is presented formally in Algorithm 2. This closure computation algorithm, introduced in Hordijk and Steel (2004), is equivalent to the “network expansion” algorithm (Ebenhöh et al., 2004) used in Goldford et al. (2017).

#### Algorithm 2 ComputeClosure (*F, R*′)

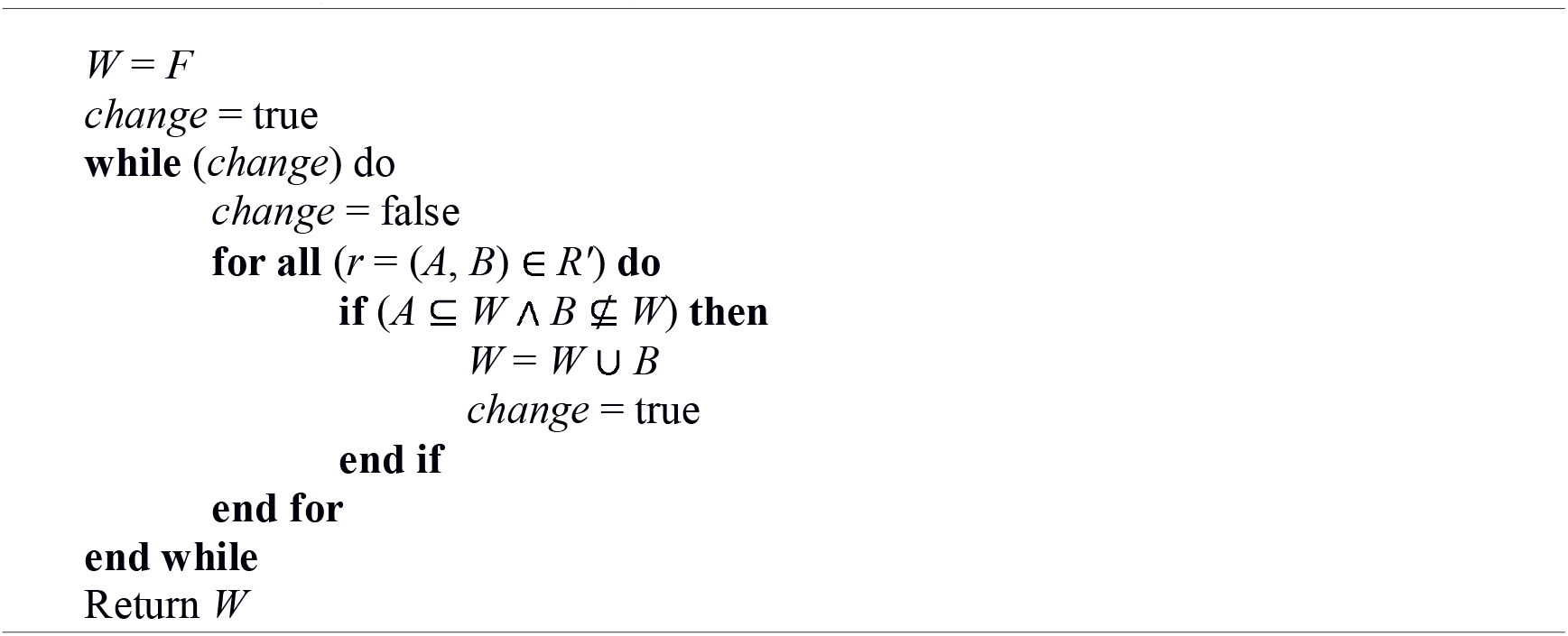

A naive computational complexity analysis of the RAF algorithm gives a worst-case running time of *O*(|*X*||*R*|^3^). However, with some additional book-keeping (such as keeping track of all reactions that each molecule is involved in), this can be reduced. In fact, the average running time on a simple polymer-based model of CRSs turns out to be sub-quadratic (Hordijk and Steel, 2004).

### LUCA enrichment analysis

The genetic families identified in Weiss et al. (2016) were mapped to KEGG orthologues, the corresponding EC numbers were retrieved and the reactions performed by these were listed and compared with the lists of reactions comprising the different networks, namely the global O_2_-independent prokaryotic network; the maxRAF obtained with this network; maxRAFs obtained with the Ace and Met subsets; and the intersection of these.

### Statistical Analysis

Fisher’s exact tests with Benjamini–Hochberg multiple test corrections were performed for pathway and cofactor enrichment analysis (Figures 3, 6A-B), and significance was considered for corrected *p*-values smaller than 0.05. A Fisher test was performed for enrichment in LUCA genes (Figure 6C) and significance was considered for *p*-values smaller than 0.0001. All statistical analysis were performed in Python ver. 3.6.6 with the package scipy.stats. Network properties were calculated and visualizations were produced with Cytoscape (Shannon, 2003) ver. 3.7.1.

### Software availability

A custom-made implementation of the maxRAF algorithm was used for the analysis in this paper and is available at https://www.canterbury.ac.nz/engineering/schools/mathematics-statistics/research/bio/downloads/raf/. An example of an input file (global prokaryotic O_2_-independent network, food set with all small molecules, abiotic carbon and organic cofactors) is given in **Data S1**. A more general-purpose and interactive RAF analyzer can be found online at http://www.math.canterbury.ac.nz/bio/RAF/, including several more examples and explanations.

## Acknowledgements

We thank Filipa L. Sousa and Martina Preiner for comments. This work was supported by grants from the European Research Council (666053) and the Volkswagen Foundation (93 046) to WFM. WH thanks the Institute for Advanced Study, University of Amsterdam, The Netherlands, for partial support in the form of a fellowship.

## Author contributions

JCX collected and integrated data from databases, curated models, and literature. JCX, MS, SK and WFM designed the project. WH and MS wrote the pseudocode and algorithm for detection of maxRAFs and WH performed maxRAF identification. JCX and WFM wrote the first manuscript draft. JCX, WH, SK, MS, and WFM contributed in data interpretation and writing the final manuscript.

## Supplementary Information Titles and Legends

**Figure S1. Pipeline of reconstructing catalysis-annotated metabolic networks.** Steps in grey include metabolic data only, steps in brown include catalysis rules, and steps in greens represent the inclusion of curated data from metabolic models of *Moorella thermoacetica* and *Methanococcus maripaludis*.

**Figure S2. MaxRAF obtained with the network of *Moorella thermoacetica*.** Node size is scaled according to the degree, with food molecules highlighted in green and relevant products in dark blue (only metabolic interconversions are depicted; catalysis arcs are omitted for clarity). ‘Acceptor’ and ‘Reduced Acceptor’ are abstract redox molecules as represented in KEGG metabolism.

**Figure S3. MaxRAF obtained with the network of *Methanococcus maripaludis*.** Node size is scaled according to the degree, with food molecules highlighted in green and relevant products in dark blue (only metabolic interconversions are depicted; catalysis arcs are omitted for clarity). ‘Acceptor’ and ‘Reduced Acceptor’ are abstract redox molecules as represented in KEGG metabolism.

**Figure S4. MaxRAF obtained with the intersection of the networks of *Methanococcus maripaludis* and *Moorella thermoacetica*.** Node size is scaled according to the degree, with food molecules highlighted in green and relevant products in dark blue (only metabolic interconversions are depicted; catalysis arcs are omitted for clarity). ‘Acceptor’ and ‘Reduced Acceptor’ are abstract redox molecules as represented in KEGG metabolism.

**Figure S5. MaxRAF obtained with amino acids and bases**. The network represents the maxRAF obtained with the full prokaryote O_2_-independent network with inorganic catalysts, abiotic compounds, all amino acids and bases but no organic cofactors added to the food set (only metabolic interconversions are depicted; catalysis arcs are omitted for clarity). Node size is scaled according to the degree, with food molecules highlighted in green. ‘Acceptor’ and ‘Reduced Acceptor’ are abstract redox molecules as represented in KEGG metabolism.

**Table S1. Metabolic networks annotated with catalysis rules. (A)** Prokaryotic, O_2_-independent global metabolic network **(B)** subset network of *Moorella thermoacetica* **(C)** subset network of *Methanococcus maripaludis*.

**Table S2. Composition of Food Sets used in predictions of maxRAFs in different metabolic networks and resulting maxRAF sizes.**

**Table S3. Lists of reactions in all maxRAFs predicted for all networks in all food sets.**

**Table S4. Connectivity of metabolites in LUCA’s maxRAF**.

**Data S1. Input file with the global prokaryotic O_2_-independent network used to run the maxRAF algorithm**. Food set with all small molecules, abiotic carbon and organic cofactors.

