## Supplementary material for "Autocatalytic chemical networks preceded proteins and RNA in evolution": Figure S1

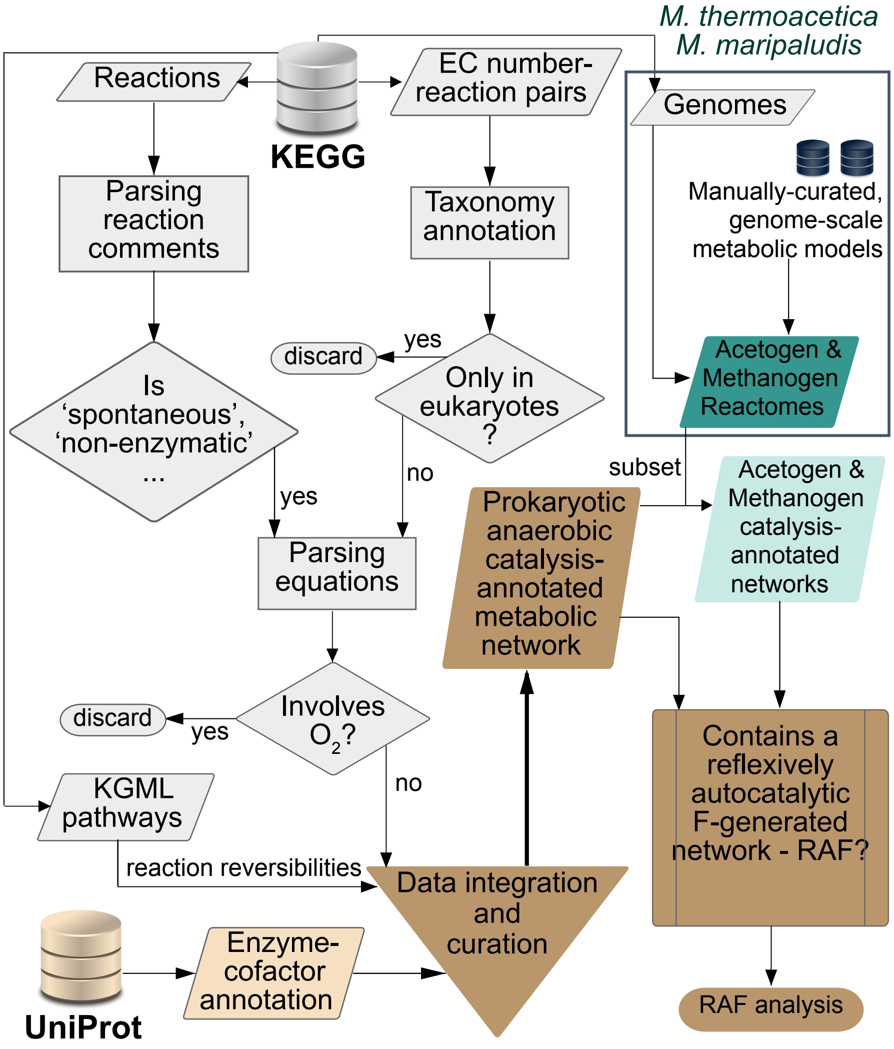


**Figure S1. Pipeline of reconstructing catalysis-annotated metabolic networks**. Steps in grey include metabolic data only, steps in brown include catalysis rules, and steps in greens repre-sent the inclusion of curated data from metabolic models of Moorella thermoacetica and Methanococcus maripaludis.
