## Supplementary material for "Autocatalytic chemical networks preceded proteins and RNA in evolution": Figure S3

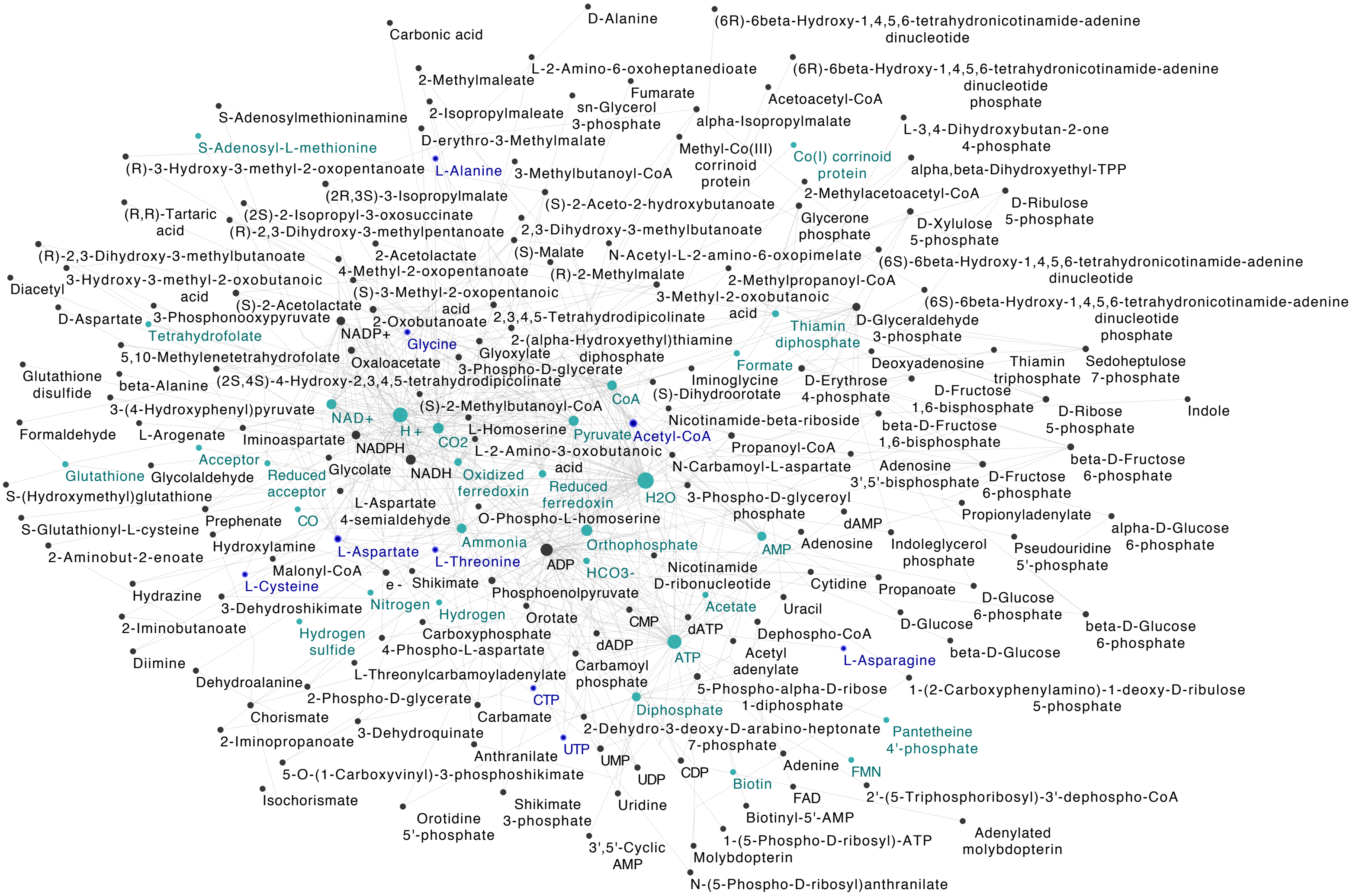


**Figure S3. MaxRAF obtained with the network of *Methanococcus maripaludis*.** Node size is scaled according to the degree, with food molecules highlighted in green and relevant prod-ucts in dark blue (only metabolic interconversions are depicted; catalysis arcs are omitted for clarity). ‘Acceptor’ and ‘Reduced Acceptor’ are abstract redox molecules as represented in KEGG metabolism.
