## Supplementary material for "Autocatalytic chemical networks preceded proteins and RNA in evolution": Figure S4

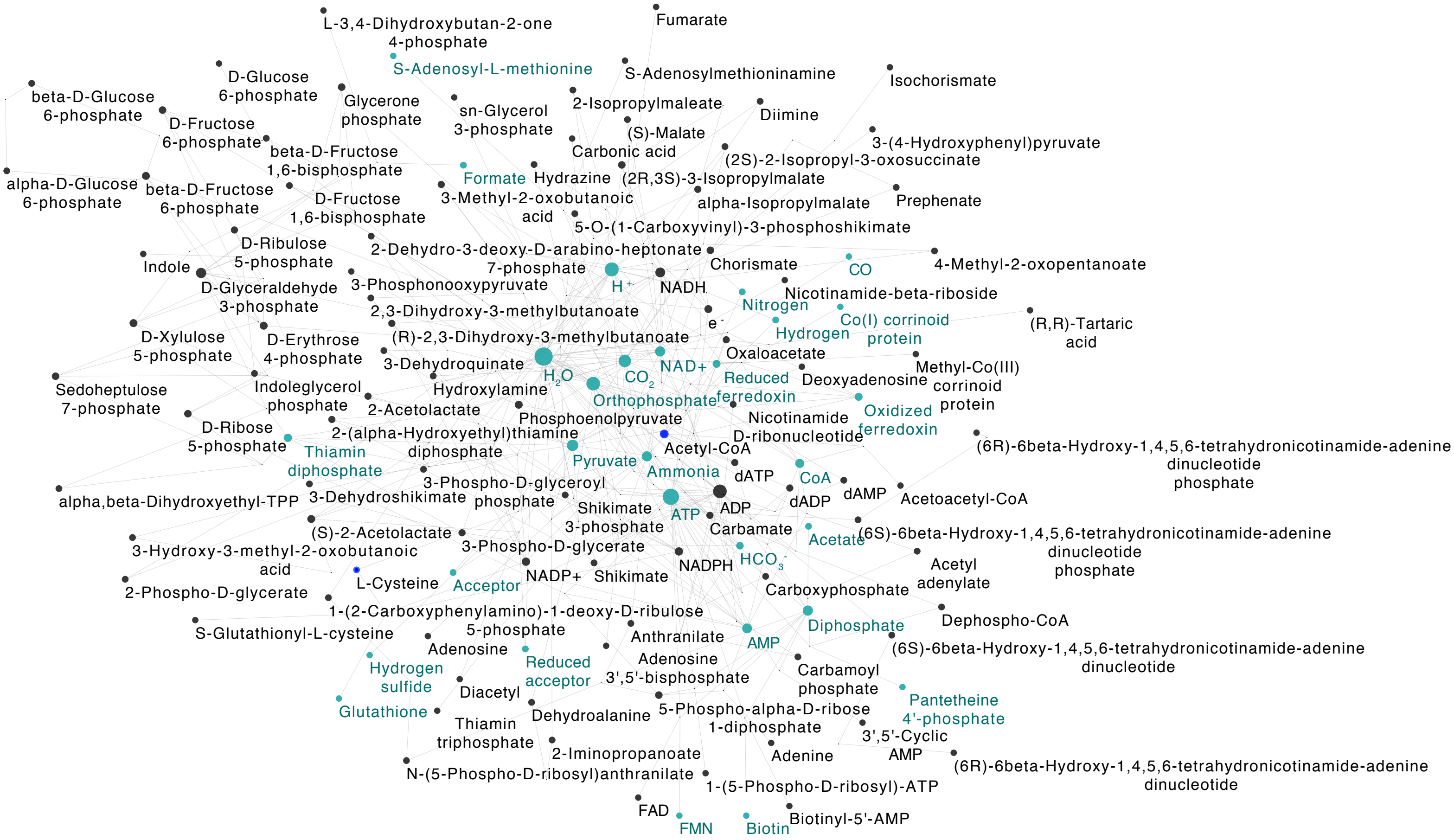


**Figure S4. MaxRAF obtained with the intersection of the networks of *Methanococcus maripaludis* and *Moorella thermoacetica*.** Node size is scaled according to the degree, with food molecules highlighted in green and relevant products in dark blue (only metabolic interconversions are depicted; catalysis arcs are omitted for clarity). ‘Acceptor’ and ‘Reduced Acceptor’ are abstract redox molecules as represented in KEGG metabolism.
