## Supplementary material for "Autocatalytic chemical networks preceded proteins and RNA in evolution": Figure S5

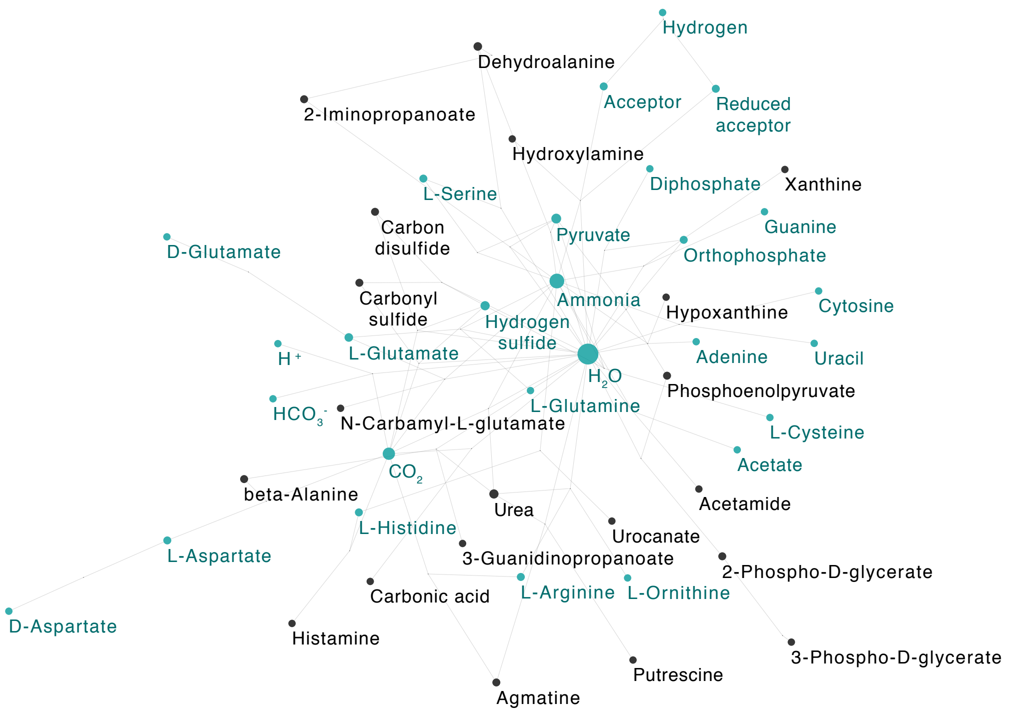


**Figure S5. MaxRAF obtained with amino acids and bases**. The network represents the maxRAF obtained with the full prokaryote O2-independent network with inorgan-ic catalysts, abiotic compounds, all amino acids and bases but no organic cofactors added to the food set (only metabolic interconversions are depicted; catalysis arcs are omitted for clarity). Node size is scaled according to the degree, with food molecules highlighted in green. ‘Acceptor’ and ‘Reduced Acceptor’ are abstract redox molecules as represented in KEGG metabolism.
