## Supplementary material for "Autocatalytic chemical networks preceded proteins and RNA in evolution": Table S2

**Table S2. Composition of Food Sets used in prediction of maxRAFs in different metabolic networks and resulting maxRAF sizes.**

| ID | Description | Size | Content | maxRAF size | | |
|---|---|---|---|---|---|---|
| | | | | Global oxygen-free network | *M. thermoacetica* | *M. maripaludis* |
| **1** | Small-molecules + inorganic catalysts | 39 | $H_2O$, $H_2$, $H^+$, $CO_2$, CO, $PO_4^{3-}$, $SO_4^{2-}$, $HCO_3^-$, $P_2O_7^{4-}$, S, $H_2S$, $NH_3$, $N_2$, CO, all metals, Fe–S clusters, Ni–Fe–S cluster, other clusters, general acceptor, general reduced acceptor (donor), general metal, "Pooling", "spontaneous" | 8 | 4 | 4 |
| **2** | **1** + abiotic organic carbon | 43 | **FS1** + Acetate, Pyruvate, Formate and Methanol | 16 | 9 | 8 |
| **3** | **2** + organic cofactors | 68 | **FS2** + FMN, Pyridoxal 5-phosphate, Thiamine diphosphate, NAD+, Molybdopterin, Cob(II)alamin, Pyrrolo-quinoline quinone, (R)-Lipoate, ATP, Biotin, Glutathione, Decylubiquinone, S-Adenosyl-L-methionine, Other quinones, Tetrahydrofolate, dipyrromethane, TTQ, 5-Hydroxybenzimidazolylcob(I)amide, AMP, Co(I) corrinoid protein, Pantetheine 4'-phosphate, Menaquinone, CoA, reduced ferredoxin, oxidized Ferredoxin | 1335 | 394 | 209 |
| **4** | **3** + Peptide | 69 | **FS3** + Peptide | 2603 | 493 | 307 |
| **5** | **2 +** aa and bases | 68 | **FS2** + all 20 amino acids, Adenine, Guanine, Cytosine, Thymine and Uracil | 33 | 19 | 14 |
